# Molecularly distinct cores coexist inside stress granules

**DOI:** 10.1101/663955

**Authors:** Luca Cirillo, Adeline Cieren, Monica Gotta

## Abstract

Stress granules are membraneless organelles that form in eukaryotic cells after stress exposure. Stress granules are constituted by a stable core and a dynamic shell that establishes a liquid-liquid phase separation with the surrounding cytosol. The structure and assembly of stress granules and how different components contribute to their formation are not fully understood. Here, using super resolution and expansion microscopy, we find that the stress granule component UBAP2L and the core protein G3BP1 occupy different domains inside stress granules. Since UBAP2L displays typical properties of a core protein, our results indicate that different cores coexist inside the same granule. Consistent with a role as a core protein, UBAP2L is required for stress granule assembly in several stress conditions and reverse genetics show that it acts upstream of G3BP1. We propose a model in which UBAP2L is an essential stress granule nucleator that facilitates G3BP1 core formation and stress granule assembly and growth.

## INTRODUCTION

Eukaryotic cells have evolved a series of mechanisms to counterbalance external stresses that may pose a threat to their internal homoeostasis. One of these mechanisms is the inhibition of protein translation and the consequent formation of membraneless organelles known as stress granules (Kedersha et al., 2000; reviewed in Mahboubi and Stochaj, 2017; Panas et al., 2016; Protter and Parker, 2016). Following translation inhibition, polysomes disassemble releasing 48S preinitiation complexes (PICs). In the cytosol, mRNA, PICs and other proteins coalesce in stress granule cores (Kedersha et al., 2000). Stress granule cores recruit a shell, whose properties are dominated by weak interactions between proteins and RNAs (Jain et al., 2016; Treeck et al., 2018; Wheeler et al., 2016). The stress granule shell establishes a liquid-liquid phase separation with the surrounding cytoplasm, explaining the high dynamicity of stress granule components (Han et al., 2012; Kato et al., 2012; Weber and Brangwynne, 2012). A mature stress granule will be formed by a phase separated shell surrounding several stable cores (Jain et al., 2016). However, the identity of the stress granule cores, their formation and localization pattern within the granules remain incompletely understood.

Proteins that contribute to the formation of stress granules include G3BP1 and 2 (Ras GTPase-activating protein-binding protein 1 and 2), TIA-1 (T-cell-restricted intracellular antigen-1), TIAR (TIA-1-related), TTP (Tristetraprolin) and FMRP (Fragile X mental retardation protein 1 homolog) (Kedersha et al., 2000; Mazroui et al., 2002; Stoecklin et al., 2004; Tourrière et al., 2003). Although the composition of stress granules as well as the requirement for specific proteins vary according to the stress-inducing agent and cell type (Aulas and Vande Velde, 2015; Bounedjah et al., 2014; Markmiller et al., 2018), G3BP1 and 2 have been shown to be required for stress granule nucleation in most cases (Kedersha et al., 2016; Matsuki et al., 2013) and G3BP1 is highly enriched in stress granule cores (Jain et al., 2016; Markmiller et al., 2018; Wheeler et al., 2016; Youn et al., 2018). The relative stability of stress granule cores has recently allowed their purification and the identification of novel stress granule nucleators, among them the protein UBAP2L (Ubiquitin-associated protein 2-like (Markmiller et al., 2018; Youn et al., 2018). UBAP2L is a ubiquitous and abundant protein in human tissues (Beck et al., 2011; Lonsdale et al., 2013; Melé et al., 2015; Uhlén et al., 2015), it promotes cancer progression and metastasis (Aucagne et al., 2017; Chai et al., 2016; He et al., 2018; Li and Huang, 2014; Li et al., 2018; Ye et al., 2017, P. 20; Zhao et al., 2015) and may be involved in neurodegenerative diseases (Markmiller et al., 2018). Despite its abundance and potential implication in diseases, the exact role of UBAP2L in cell physiology and stress granule formation is not yet understood. Here, we used a combination of cell biology and reverse genetics to investigate the role of UBAP2L is stress granule formation. We showed that UBAP2L is present in discrete particles inside stress granules, which do not colocalize with G3BP1 stress granule cores. UBAP2L acts upstream of G3BP1 and 2 in stress granule assembly but does not interfere with stress-induced translational inhibition, indicating that UBAP2L is a crucial stress granule nucleator downstream of translational inhibition. Our results show that molecularly distinct stress granule cores coexist in the same stress granules.

## RESULTS

### UBAP2L localizes inside stress granules in discrete particles distinct from G3BP1 cores

Given the importance of UBAP2L in sodium arsenite (arsenite hereafter)-induced stress granule assembly (Markmiller et al., 2018; Youn et al., 2018), we sought to investigate its sublocalization inside stress granules. To this end, we used 3D-STED (3D Stimulated Emission Depletion miscroscopy, Hell and Wichmann, 1994; Klar and Hell, 1999). 3D-STED immunofluorescence of HeLa K cells fixed and stained with anti-UBAP2L antibodies, revealed that each stress granule is formed by discrete UBAP2L particles of roughly round shape (Fig. 1 A), with a diameter of 187.0 ± 44.2 nm (mean ± SD, Fig. 1 B). We also performed 3D-STED on HeLa K cells stained for the known stress granule component G3BP1 and measured the diameter of G3BP1 cores (Fig. 1 A). Our analysis revealed a diameter of 182.7 ± 41.9 nm (mean ± SD), which is in good agreement with the published diameter of 190.8 ± 28.3 nm (Fig. 1B, Jain et al., 2016). The number of UBAP2L and G3BP1 particles linearly correlated with the stress granule volume (Fig. S1 A). The localization of UBAP2L in discrete particles and their size distribution suggest that UBAP2L localizes to stress granule cores.

**Figure 1.**
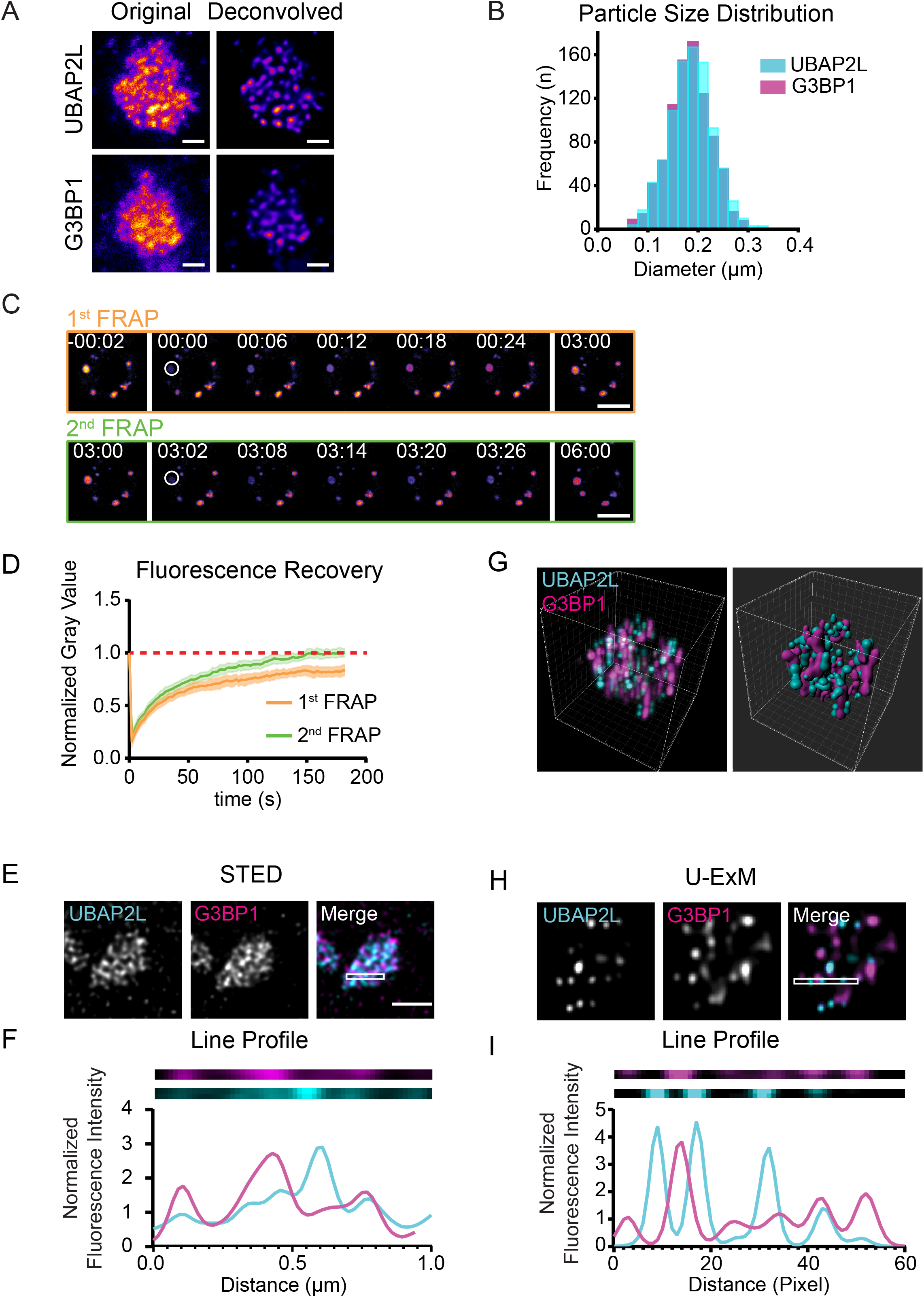
Inside stress granules UBAP2L forms discrete particles that do not colocalize with G3BP cores. A) 3D-STED images of a stress granule (detected by anti-UBAP2L or anti-G3BP staining) in HeLa K cells treated with 0.1 mM arsenite for 30 minutes prior to fixation. Scale bars correspond to 0.5 µm. B) Size distribution of stress granule particles of UBAP2L and G3BP. C) Representative images over time of a FRAP experiment on UBAP2L-mScarlet HeLa cells treated with 0.5 mM arsenite. White circle represents the bleached area. Time is expressed at mm:ss and indicated on the image, top left. Scale bars correspond to 10 µm. The red dotted line centered at 1 highlights complete fluorescence recovery. D) Average FRAP recovery curves of the first FRAP cycle and the second FRAP cycle. Mean ± 95% Confidence Interval (C.I.), n= 27 stress granules from 27 cells. E) Deconvolved 2D-STED images of a stress granule in HeLa K cells treated with 0.1 mM arsenite. Cells were stained with anti-UBAP2L and anti-G3BP antibodies. Scale bar corresponds to 1 µm. F) Line profile of the region of interest (ROI) highlighted by the white boxes in (E). The two top panels represent the magnification of the ROI split into two channels: UBAP2L (Cyan) and G3BP (Magenta). The graph corresponds to the normalized pixel-by-pixel gray value of the two images above. G) Left: Deconvolved 3D confocal image of U-ExM of a stress granule in HeLa K cells treated 30 minutes with 0.1 mM arsenite and stained with anti-UBAP2L and anti-G3BP antibodies. Right: 3D reconstruction of the surface of UBAP2L (Cyan) and G3BP (Magenta) particles. H) Deconvolved 3D confocal image of U-ExM of a stress granule in HeLa K cells treated 30 minutes with 0.1 mM arsenite, stained with anti-UBAP2L and anti-G3BP antibodies. I) Line profile of the ROI highlighted by the white boxes in (G). The two top panels represent the magnification of the ROI split into two channels: UBAP2L (Cyan) and G3BP (Magenta). The graph corresponds to the normalized pixel-by-pixel gray value of the two images above.

The shell and the core of stress granules do not only differ in their composition, but also in the dynamics of their components, with core proteins being more stable and shell proteins more dynamic. To assess whether UBAP2L particles are stress granule cores, we measured the dynamics of UBAP2L in living cells. To visualize UBAP2L, we tagged endogenous UBAP2L at the C-terminus with the red fluorescent protein mScarlet using CRISPR-Cas9 (Fig. S1 B Bindels et al., 2017). Clones were screened via PCR (Polymerase Chain Reaction, Fig. S1 C) and validated with western blot using an anti-UBAP2L antibody (Fig. S1 D). In addition, treatment of UBAP2L-mScarlet HeLa cells with a siRNA directed against mScarlet, followed by western blot analysis, confirmed the successful insertion of the construct (Fig. S1 E). Although we failed to identify a homozygous insertion (Fig. S1 D, E), live cell imaging revealed that UBAP2L-mScarlet transitions from a diffuse cytosolic signal to a droplet-like pattern following arsenite exposure (Fig. S1 F). This observation is consistent with published immunofluorescence of UBAP2L (Markmiller et al., 2018; Youn et al., 2018), indicating that UBAP2L-mScarlet recapitulates the localization of wild-type UBAP2L. Having created a tool to visualize UBAP2L *in vivo*, we next measured the dynamics of UBAP2L in stress granules using Fluorescence Recovery after Photobleaching (FRAP). After treating UBAP2L-mScarlet HeLa cells with arsenite to induce stress granule formation, we bleached a single cytosolic granule and followed the fluorescence recovery over time using confocal microscopy (Fig. 1 C). Analysis of fluorescence recovery for 3 minutes after bleaching, highlighted the presence of a slow recovery fraction of UBAP2L-mScarlet (Immobile Fraction = 18.2 ± 1.9 %, mean ± SD, Fig. 1 D, Fig. S1 G, H). To validate the presence of a slow recovery fraction we repeated the bleaching cycle in the same stress granule in the context of a double FRAP experiment (Krouwels et al., 2005; Stavreva and McNally, 2004). If the first photobleaching event bleaches all the fluorophores of the immobile fraction, the recovery curve after the second bleaching of the same granule depends uniquely on the mobile fraction and should yield a 100% recovery. We obtained a complete recovery of the fluorescence 3 minutes after the second bleaching (Immobile Fraction = −0.05 ± 2.1 %, mean ± SD, Fig. 1 D, Fig. S1 G, H). Moreover, the second recovery curve is significantly different from the first recovery curve (p<0.0001, two-way ANOVA). This experiment demonstrates that UBAP2L-mScarlet exists in stress granules in at least two different pools, characterized by different diffusivity, in agreement with previous data on other stress granule proteins (Jain et al., 2016) and mRNAs (Mollet et al., 2008; van der Laan et al., 2012).

The localization in discrete particles and the presence of an immobile fraction suggest that UBAP2L behaves as stress granule core protein. We therefore, asked whether UBAP2L particles colocalize with G3BP1. We performed two-colors 2D STED on arsenite-treated HeLa K cells fixed and stained with both anti-UBAP2L and anti-G3BP antibodies. Contrary to our expectations, most UBAP2L and G3BP1 stress granule particles did not colocalize (Fig. 1 E, F, Pearson correlation coefficient: 0.01 ± 0.25, mean ± SD). To confirm these observations, we investigated the localization of UBAP2L with an independent approach. Ultrastructure Expansion Microscopy (U-ExM) allows imaging beyond the resolution limits of visible light by expanding the sample without affecting its ultrastructure (Gambarotto et al., 2019). We performed U-ExM on arsenite-induced stress granules in Hela K cells fixed and stained with antibodies against UBAP2L and G3BP1. With an estimated expansion between 3.5 and 4.0-fold compared to unexpanded samples, three-dimensional confocal microscopy revealed that UBAP2L and G3BP1 localize in discrete patterns inside stress granules (Fig. 1 G, H, I), supporting our 3D-STED results (Fig. 1 A). In addition, in agreement with 2D-STED (Fig. 1 E, F), only a minority of UBAP2L and G3BP particles colocalized (Pearson Coefficient: 0.03 ± 0.19, mean ± SD).

These results demonstrate that stress granules contain at least two distinct populations of UBAP2L (a mobile fraction and an immobile one) and that UBAP2L localizes in discrete particles reminiscent of stress granule cores. However, UBAP2L particles and G3BP1 cores do not colocalize, suggesting that UBAP2L particles are distinct from G3BP1 stress granule cores.

### UBAP2L is essential for stress granule assembly induced by many cellular stressors

G3BP1 forms particles inside stress granules (Jain et al., 2016) and is an important regulator of stress granule formation (Kedersha et al., 2016). Our data demonstrate that UBAP2L also exists in particles, suggesting it may play an important role in stress granule assembly, like G3BP1.

Since stress granules induced by different stressors show different compositions (Fujimura et al., 2012; Kedersha et al., 2008), we asked whether UBAP2L is a general stress granule component or whether it displays a stress-specific recruitment. We exposed HeLa K cells to a variety of stresses (see Material and Methods) and stained them with anti-G3BP antibodies as a marker for stress granules and with anti-UBAP2L antibodies. We imaged the cells with confocal microscopy and evaluated the Pearson colocalization coefficient of UBAP2L with G3BP1 (Fig. 2A, B). We found that UBAP2L formed granules that colocalized with G3BP1 stress granules in arsenite treatment, endoplasmic reticulum (ER) stress (DTT), proteasome inhibition (MG132), heat shock and mitochondrial stress (NaN_3_, Fig 2 A, B). The colocalization score was low after the 90° rotation in the cytoplasm of non-treated control cells and in arsenite-treated cells, excluding the possibility of a random colocalization between the two fluorescent signals (Fig. S2A, B).These results show that UBAP2L is a general component of stress granules, consistent with recent data (Huang et al., 2019).

UBAP2L is required for stress granule assembly in cells stressed with arsenite, H_2_O_2_ and Sorbitol (Huang et al., 2019; Markmiller et al., 2018; Youn et al., 2018). We next tested whether UBAP2L was also required for stress granule assembly in the conditions that we tested above (ER stress, proteasome inhibition, heat shock and mitochondrial stress). We depleted UBAP2L using two independent siRNAs and found that both siRNAs strongly reduce UBAP2L signal (97.0 ± 1.9 % depletion for siUBAP2L #1, 97.6 ± 1.7 % depletion for siUBAP2L #2, mean ± SEM) when compared to cells transfected with a control siRNA (Fig. S2 C, D). We then visualized stress granules with anti-G3BP antibodies in cells exposed to ER and mitochondrial stress, proteasome inhibition and heat shock in the absence of UBAP2L. In control-depleted cells, stress granules promptly assembled following stress, whereas non-treated cells rarely assembled stress granules (Fig. 2 C, D). Different treatments result in the formation of stress granules of different size and shape (Fig. 2 C), as previously described (reviewed in Mahboubi and Stochaj, 2017; Panas et al., 2016; Protter and Parker, 2016). Compared to control depleted cells, UBAP2L depletion cause a significant reduction of the percentage of cells displaying G3BP1-positive stress granules (Fig. 2 C, D). We observed the same requirement for UBAP2L in arsenite-induced stress granule formation when using antibodies against TIA-1 or USP10 as alternative stress granule markers, excluding that the role of UBAP2L is limited to G3BP1 (Fig. S2 E).

These results show that UBAP2L is a general component of stress granules and it is not only required for arsenite-induced stress granule assembly (Markmiller et al., 2018; Youn et al., 2018) but also for stress granules assembly in other stress conditions, consistent with recent data (Huang et al., 2019). It is interesting that UBAP2L is required for stress granule assembly in heat-shock conditions. G3BP1 is not required for heat-induced stress granule formation (Kedersha et al., 2016), suggesting that UBAP2L plays a more general role than G3BP1.

**Figure 2.**
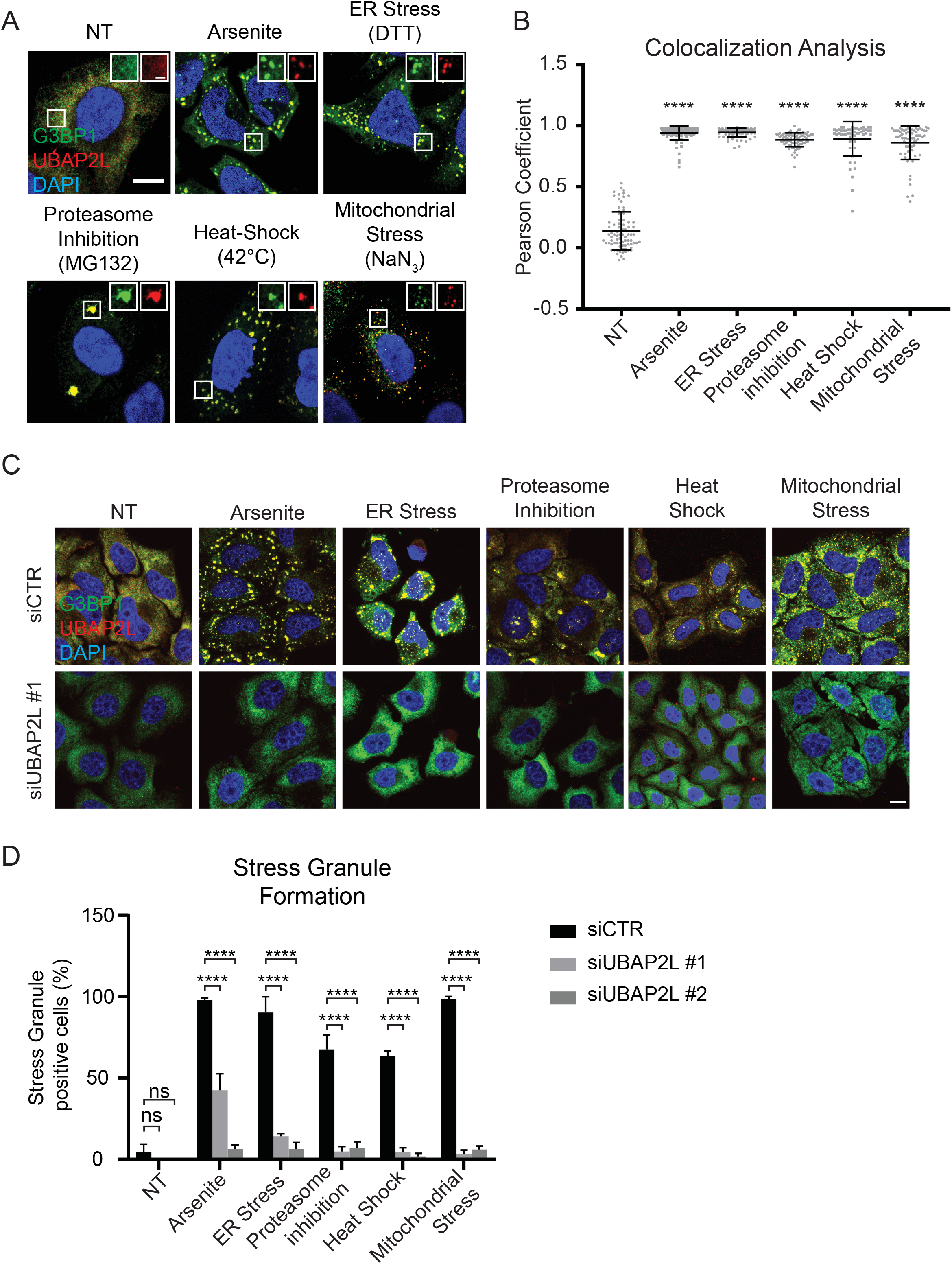
UBAP2L localizes to and is required for stress granule formation after exposure to diverse stress conditions. A) Confocal images of HeLa K cells stained with anti-G3BP and anti-UBAP2L antibodies and counterstained with DAPI. In all figures, NT is non-treated cells. White box indicates the ROI enlarged and depicted in separate channels on the top right. The scale bar in the main image corresponds to 10 µm, the scale bar in the ROI corresponds to 1 µm. B) Scatterplot representing the cross correlation between G3BP1 and UBAP2L signal. Error bars indicate SD. Statistical significance was determined using ANOVA and Dunnett’s multiple comparison. Each dot corresponds to the Pearson coefficient of a single ROI. N = 3 independent experiments. In all figures and supplementary figures, ns, p > 0.05, *p < 0.05, **p < 0.01, ***p < 0.001, ****p < 0.0001. C) Maximum projections of confocal images of HeLa K cells transfected with the indicated siRNAs, stained with anti-G3BP and anti-UBAP2L antibodies and counterstained with DAPI. Scale bar corresponds to 10 µm. D) Percentage of HeLa K cells transfected with the indicated siRNAs containing G3BP1-positive stress granules. Error bars indicate Standard Error of the Mean (SEM). Statistical significance was determined using two-way ANOVA and Tukey’s multiple comparison. N ≥ 2 independent experiments.

### UBAP2L promotes stress granule assembly upstream of G3BP1

Stress granules cannot form in absence of the proteins G3BP1 and G3BP2 (Kedersha et al., 2016; Matsuki et al., 2013). To test whether UBAP2L requires G3BP1 and 2 for its localization to stress granules, we co-depleted both G3BP1 and 2 in HeLa K cells (Fig. S3). Following UBAP2L depletion, we induced stress granule formation using arsenite and then fixed and stained cells with anti-UBAP2L and anti-G3BP antibodies (Fig 3 A). Compared to control-depleted cells, the percentage of cells containing UBAP2L positive granules did not significantly change when G3BP1 and 2 were depleted (Fig. 3 B), suggesting that G3BP1 and 2 are dispensable for stress granules recruitment of UBAP2L.

**Figure 3.**
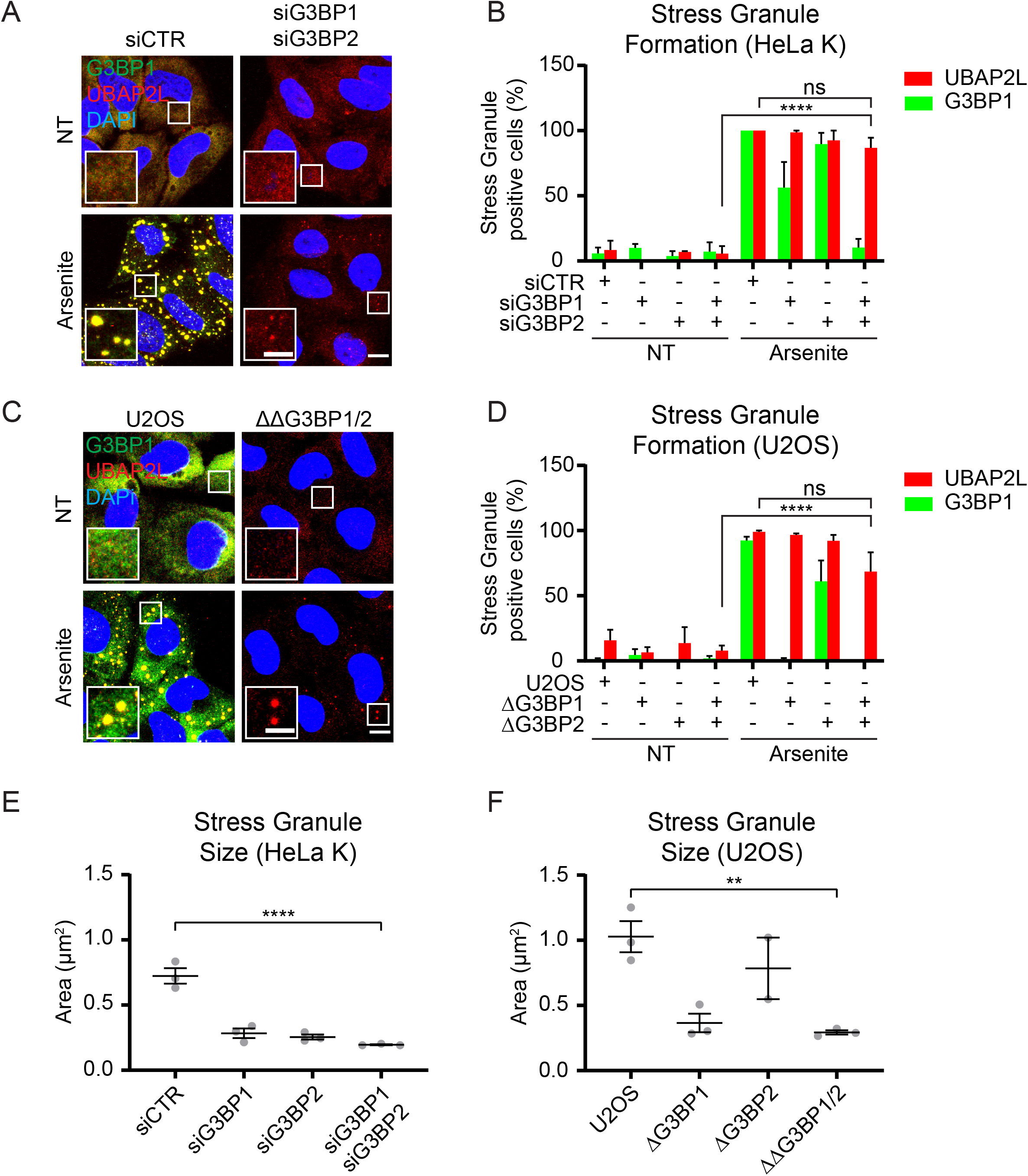
Recruitment of UBAP2L to stress granules is independent of G3BP. A) Maximum projections of confocal images of HeLa K cells transfected with the indicated siRNAs and stained with anti-G3BP, anti-UBAP2L antibodies and counterstained with DAPI. Cells were either left untreated (upper panels), treated with 0.1 mM arsenite for 30 minutes (lower panels). White box indicates the ROI endlarged and depicted in separate channels on the bottom left. Scale bar in the image corresponds to 10 µm, scale bar in the ROI corresponds to 2 µm. B) Percentage of HeLa cells transfected with the indicated siRNA, containing either UBAP2L positive stress granules or G3BP1 positive stress granules. Cells were treated with arsenite 0.1 mM or left untreated. N≥2 independent experiments. Statistical significance was determined using two-way ANOVA and Tukey’s multiple comparison. C) Maximum projections of confocal images of U2OS and U2OS ΔΔG3BP1/2 cells stained with anti-G3BP and anti-UBAP2L antibodies and counterstained with DAPI. Cells were either left untreated (top) or treated 60 minutes with 0.1 mM arsenite (bottom). White box highlights the ROI corresponding to the detail enlarged in the bottom left of the image. Scale bar in the image corresponds to 10 µm. Scale bar in the detail corresponds to 2 µm. D) Percentage of U2OS, U2OS ΔG3BP1, U2OS ΔG3BP2, U2OS ΔΔG3BP1/2 cells containing either UBAP2L positive stress granules or G3BP1 positive stress granules after 60 minutes exposure to 0.1 mM arsenite. N=3 independent experiments. Statistical significance was determined using two-way ANOVA and Tukey’s multiple comparison. E) Quantification of UBAP2L-positive stress granule areas in HeLa K cells transfected with the indicated siRNA and treated 30 minutes with 0.1 mM arsenite. Mean ± SEM. Each dot represents an independent experiment. Statistical significance was determined using two-way ANOVA and Dunnett’s multiple comparison. F) Quantification of UBAP2L-positive stress granule areas in U2OS, U2OS ΔG3BP1, U2OS ΔG3BP2, U2OS ΔΔG3BP1/2 cells, 60 minutes after 0.1 mM arsenite treatment. Mean ± SEM. Each dot represents an independent experiment. Statistical significance was determined using two-way ANOVA and Dunnett’s multiple comparison.

Because of the pitfalls of transient protein depletion, we cannot formally exclude that a small pool of G3BP1 and 2 is sufficient for the formation of UBAP2L positive stress granules. To tackle this problem, we used G3BP1 and 2 knock-out U2OS cells (Fig. 3 C, Kedersha et al., 2016). Immunofluorescence analysis of these cells after arsenite treatment revealed that UBAP2L localizes into cytosolic granules (Fig 3 C, D), confirming that UBAP2L granule formation is independent of G3BP1 and 2. In both HeLa K and U2OS cell lines the depletion or the knock-out of G3BP1 and 2 result in a reduction of stress granule size compared to control cells (Fig. E, F) indicating that G3BP1 and 2 are important for stress granule maturation.

Our results support a model in which UBAP2L acts upstream of other stress granule nucleators such as G3BP1 and 2.

### Inhibition of translation after stress does not depend on UBAP2L

Since our results show that UBAP2L is necessary for stress granule assembly in several conditions and acts upstream of G3BP1, we reasoned that the absence of UBAP2L may affect the signaling pathway upstream of stress granule nucleation. Stress granule assembly is a multistep process that requires inhibition of translation and polysome disassembly (reviewed in Mahboubi and Stochaj, 2017; Panas et al., 2016; Protter and Parker, 2016). An early step in arsenite-induced stress granule assembly is the phosphorylation of the elongation factor eIF2α (Eukaryotic Initiation Factor 2 α) on Serine 51. To assess whether UBAP2L contributes to eIF2α phosphorylation in stress conditions, we depleted UBAP2L and exposed the cells to arsenite. Western blot using an antibody directed against pS51-eIF2α revealed no difference between cell lysates depleted for UBAP2L and control (Fig. 4 A), indicating that UBAP2L it is not required for eIF2α phosphorylation after stress.

**Figure 4.**
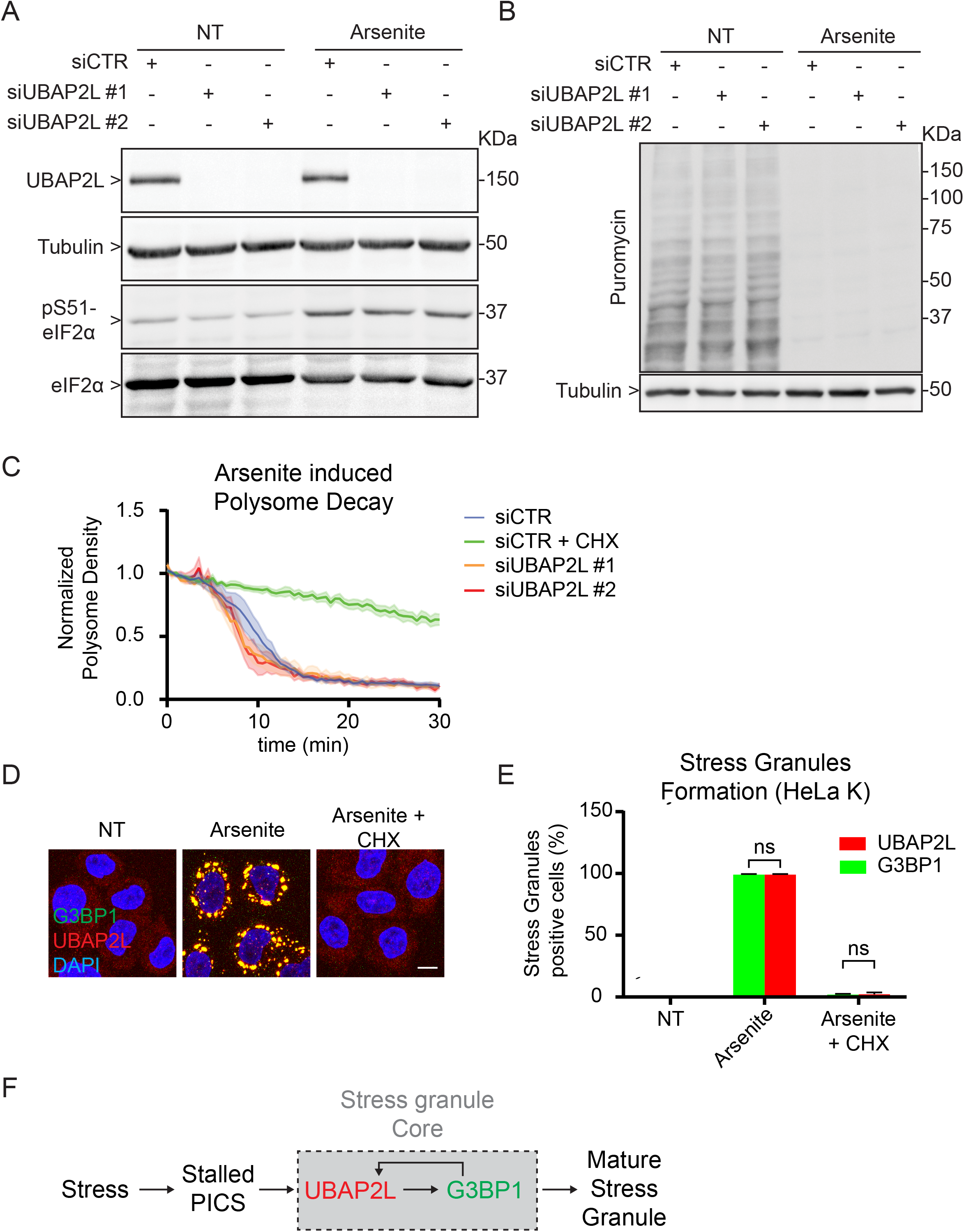
UBAP2L depletion does not affect stress induced translational inhibition. A) Representative image of a Western blot of lysates from HeLa K cells transfected with a control siRNA (1^st^ and 4^th^ lane) compared to lysates of HeLa K transfected with siRNAs targeting UBAP2L (2^nd^, 3^rd^, 5^th^ and 6^th^ lane). Cells where either left untreated (1^st^, 2^nd^, 3^rd^ lane) or treated 30 minutes with 0.1 mM arsenite (4^th^, 5^th^, 6^th^ lane). The membrane was blotted with anti-UBAP2L, anti-αTubulin, anti-pS51-eIF2α and anti-eIF2α antibodies, as indicated. N=3 independent experiments. B) Representative image of a Western blot of lysates from HeLa K cells transfected with a control siRNA (1^st^ and 4^th^ lane) compared to lysates of HeLa K transfected with siRNAs targeting UBAP2L (2^nd^, 3^rd^, 5^th^ and 6^th^ lane). Cells where either left untreated (1^st^, 2^nd^, 3^rd^ lane) or treated 30 minutes with 0.1 mM arsenite (4^th^, 5^th^, 6^th^ lane). 10 minutes before lysis cells were incubated with medium supplemented with 10 µg/ml Puromycin. The membrane was blotted with anti-Puromycin and anti-αTubulin as indicated. N=3 independent experiments. C) Quantification of a polysome decay assay. It is represented the mean normalized polysome density ± SEM over time following addition of 0.1 mM arsenite, or 0.1 mM arsenite and 50 µg/ml Cycloheximide in the culture medium. N=7 independent experiments. D) Maximum projections of confocal images of HeLa K cells stained with anti-G3BP and anti-UBAP2L antibodies and counterstained with DAPI. Cells were either left untreated (left panel), treated with 0.1 mM arsenite for 30 minutes (middle panel) or treated with both arsenite 0.1 mM and cycloheximide 50 µg/ml for 30 minutes (right panel). Scale bar in the image corresponds to 10 µm. E) Percentage of HeLa cells containing either UBAP2L positive stress granules or G3BP1 positive stress granules in untreated condition, after 30 minutes exposure to arsenite 0.1 mM, or to arsenite 0.1 mM and cycloheximide 50 µg/ml. N=3 independent experiments. F) Schematic drawing of the proposed mechanism for stress granules core formation.

Next, we asked whether cells depleted for UBAP2L efficiently inhibit translation after stress. To answer this question, we used a puromycilation assay; as puromycin covalently binds to nascent peptides, puromycin incorporation can be used as a readout for the rate of translation (Azzam and Algranati, 1973; Goodman et al., 2012; Schmidt et al., 2009). Western blot analysis with an anti-puromycin antibody revealed that arsenite treatment strongly reduces puromycin incorporation in both control and UBAP2L depleted cells (Fig. 4 B), indicating that UBAP2L is not required for arsenite induced translation inhibition.

Following translation inhibition, a crucial step in stress granule formation is the disassembly of polysomes. Drug treatments stabilizing polysomes, such as cycloheximide, inhibit stress granule formation (Bounedjah et al., 2014; Buchan et al., 2011; Kedersha et al., 2000; Mokas et al., 2009). Since we did not detect a role of UBAP2L in arsenite induced translation inhibition, we investigated the requirement of UBAP2L in polysome disassembly. To this end, we developed an assay to follow polysome disassembly in live cell imaging, using a previously published polysome reporter cell line (Pichon et al., 2016). Following siRNAs transfection, HeLa cells containing a fluorescent polysome reporter were treated with arsenite and imaged for 30 minutes. Counting the number of polysomes over time, we found that polysomes disappeared ≈15 minutes after the addition of stress in control depleted cells (Fig. 4 C). As a positive control, we compared arsenite-treated cells with cells treated with both arsenite and cycloheximide (CHX). Cells receiving this double treatment displayed only a modest rate of polysome decay, compared with cells receiving arsenite alone, indicating that our assay efficiently detects a polysome disassembly defect *in vivo* (Fig. 4 C). Similar to control depleted cells, UBAP2L depletion did not prevent arsenite-induced polysome disassembly (p=0.4692 for siUBAP2L #1 and p=0.4544 for siUBAP2L #2, two-way ANOVA and Dunnett’s multiple comparison). This result suggests that UBAP2L is not required for polysome disassembly following an arsenite treatment.

An early event in stress granule formation is the coalescence of PICs in stress granule cores (reviewed in Panas et al., 2016). UBAP2L could potentially act upstream of this process, constituting the first stress granule seed. To test this hypothesis, we stressed HeLa cells with arsenite alone or a combination of arsenite and cycloheximide. We then fixed and stained the cells with anti-UBAP2L and anti-G3BP antibodies. Following arsenite treatment, both UBAP2L and G3BP1 coalesce into stress granules (Fig. 4 D, Markmiller et al., 2018; Youn et al., 2018), whereas treatment with arsenite and cycloheximide together prevented the formation of both UBAP2L and G3BP1 positive stress granules (Fig. 4 D, E). This result shows that UBAP2L cannot nucleate stress granules before polysome disassembly and polysome-derived PIC formation.

These data together show that UBAP2L recruitment to stress granules depends on polysome derived PICs but it is independent of other stress granule nucleators.

## DISCUSSION

The inner structure of stress granules and several aspects of the pathway leading to stress granule formation remain controversial. Here, we investigated the stress granule protein UBAP2L and found that it behaves as a stress granule core component, yet it does not colocalize with G3BP1 cores. We showed that recruitment of UBAP2L to stress granules requires polysome disassembly but is independent of G3BP1, suggesting that UBAP2L coalescence precedes G3BP1 recruitment to stress granules.

Our data are not contradictory with the proposed function of G3BP1 and 2 as essential stress granule nucleators (Kedersha et al., 2016; Matsuki et al., 2013). Albeit G3BP1 and 2 are not required for the formation of UBAP2L positive stress granules, they are important to recruit other components and to form a mature stress granule (Kedersha et al., 2016; Matsuki et al., 2013). In agreement, we observed a decrease in UBAP2L granule size after G3BP1 and 2 depletion (Fig. 3 E, F). The absence of both G3BP1 and 2 results in UBAP2L positive stress granule of similar size in HeLa K (0.196 ± 0.100 µm^2^) and U2OS cells (0.283 ± 0.169µm^2^; Mean ± SD), suggesting that stress granule may have a lower size limit. These measurements are too close to the diffraction limit of our experiment to be conclusive. Nevertheless, one can speculate that in absence of G3BP1 and 2, only isolated UBAP2L cores assemble, suggesting that, in normal conditions, UBAP2L cores come first in stress granules nucleation, whereas G3BP1 cores are required later for stress granules maturation (Fig. 4 F). The upstream role of UBAP2L is also consistent with recent work showing that depletion of UBAP2L results in the loss of the interaction of G3BP2 with ribosomal subunits (Huang et al., 2019).

Altogether, our results provide evidence that core particles of different nature coexist in the same stress granule. We propose a model in which both G3BP1 and UBAP2L bind stalled preinitiation complexes, but they contribute differently to stress granule assembly (Fig. 4 F). UBAP2L acts upstream of G3BP1, preceding the formation and/or the recruitment of G3BP1 cores inside the stress granules. An intriguing possibility is that UBAP2L and G3BP1 particles reflect distinct pools of preinitiation complexes, possibly linked to different mRNA species. Further work is needed to test these hypotheses and develop a comprehensive model of stress granule nucleation. Other stress granule components, such as PABP1 and mRNAs, form a core-like pattern inside stress granules suggesting that even more core species exist (Jain et al., 2016; Souquere et al., 2009; Wheeler et al., 2016).

The role of UBAP2L in stress granule nucleation is of interest from a clinical perspective. Several stress granule proteins contribute to neurodegenerative disorders such as Amyotrophic Lateral Sclerosis (ALS) or Frontal Temporal Lobar Dementia (FTLD) (Jain et al., 2016; Markmiller et al., 2018; Youn et al., 2018; reviewed in Li et al., 2013; Ramaswami et al., 2013). For example, stress granule independent phase separation of TDP-43 (TAR) DNA binding protein), a hallmark of ALS, causes cell death in neurons (Gasset-Rosa et al., 2019), suggesting that a functional stress granule machinery has a protective effect towards neurodegeneration. In *D. melanogaster*, modulating the levels of the proposed ortholog of UBAP2L, Lingerer, has an influence on FUS and FTD/ALS-associated poly(GR) protein toxicity (Markmiller et al., 2018).

Aside from neurodegeneration, a number of publications show a role of UBAP2L in cancer. UBAP2L appears to act in different malignances as an oncogene, whose expression is required for survival (Chai et al., 2016), growth (He et al., 2018; Li and Huang, 2014; Li et al., 2018; Zhao et al., 2015), and metastasization (Aucagne et al., 2017; Li and Huang, 2014; Li et al., 2018; Ye et al., 2017). Stress granules are often present in cancer cells (reviewed in Mahboubi and Stochaj, 2017), where they may promote cell survival in the stressful condition of the cancer microenvironment (Adjibade et al., 2015; Somasekharan et al., 2015; Vilas-Boas et al., 2016) and metastasization (Somasekharan et al., 2015). Interfering with stress granule formation by reducing G3BP1 expression, has a protective effect towards metastasization in mice (Somasekharan et al., 2015). In analogy with G3BP1, inhibiting stress granule formation in cancer cells by targeting UBAP2L represents a therapeutic option that is worth further investigation.

In conclusion, our results show that UBAP2L is a core component of stress granules and is a crucial regulator of stress granule assembly in many stress conditions. UBAP2L may therefore represent an attractive therapeutic target for the treatment of neurodegenerative diseases and cancer.

## STAR METHODS

### Key Resource Table

Table S1

### Contact for Reagent and Resource Sharing

Further information and requests for resources and reagents should be directed to and will be fulfilled by the Lead Contact, Monica Gotta

### Experimental Model and Subject Details

HeLa Kyoto (HeLa K) and U2OS cells were grown in DMEM (Gibco) supplemented with 10% fetal calf serum (FCS; Gibco), 2 mM L-glutamine, 100 U/ml penicillin, and 100 μg/ml streptomycin (Invitrogen). Cells were maintained in an humidified incubator at 37°C and 5% CO_2_ concentration. For live cell imaging experiment and heat shock treatment (Fig. 1 C, S1 F, Fig. 2, Fig. 4 C) cells were imaged in Leibovitz L-15 (Thermofisher) medium supplemented with 10% FCS.

## METHOD DETAILS

### Cell transfection and treatments

For STED (Fig. 1 A, B, E, F and Fig S1 A), 300’000 cells were seeded on 20 mm high precision coverslips (Karl Hecht) in a 6 well plate/35 mm dish, 24 hours before the experiment. For experiments in Fig. 1 G, H, I and Fig. 2 and Fig. 3 cells were seeded 30-40% confluent on a 1 cm round coverslip, 24 hours prior to transfection. For experiments in Fig. 4 A, B, Fig. S1 D, E and Fig S3, 150’000 HeLa K cells were seeded in a 6 well plate/35 mm dishes, 24 hours prior to transfection. In Fig. 3C, 5000 PolII-SunTagx56-Ki67-MS2×132 HeLa cells (Pichon et al., 2016), were seeded in a 8-well IBIDI plate (Vitaris). For experiments in Fig. 4 D, E 100’000 cells were seeded on 20 mm square coverslips in a 6 well plate/35 mm dishes, 24 hours prior to transfection.

Transfection was performed with a final concentration of 20 nM siRNAs and carried out using LipofectamineTM RNAiMAX transfection Reagent according to the manufacturer’s instructions (Invitrogen). The following target sequences were used (Dharmacon):

- 5’-GCAAUAGCAGCGGCAAUACGUdTdT-3’ (siUBAP2L #1),
- 5’-CAACACAGCAGCACGUUAUdTdT-3’ (siUBAP2L #2),
- 5’- CUUACAACGUCGACCGCAAdTdT-3’ (simScarlet),
- 5’- ACAUUUAGAGGAGCCUGUUGCUGAAdTdT-3’(siG3BP1),
- 5’- GAAUAAAGCUCCGGAAUAUdTdT-3’ (siG3BP2),
- 5’- GGACCUGGAGGUCUGCUGUdTdT-3’(siCTR).

In codepletion experiments (Fig. 3 A, B, E) a total of 40 nM siRNA was used: in the single depletions 20 nM siG3BP1 or 2 were mixed with 20 nM siCTR to account for siRNA competition. In all the experiments a 48-hours siRNA treatment was performed.

For ER stress (Fig. 2), HeLa K were treated with 2 mM Dithiothreitol (DTT, Sigma) diluted into culture medium for 30 minutes at 37°C. For Proteasome Inhibition (Fig. 2), HeLa K were treated with 10 µM MG132 (Sigma) diluted into culture medium at 37°C. For heat shock (Fig. 2), HeLa K were transferred to L-15 medium and incubated for 3 hours at 37°C, then the temperature was increased at 42°C for 60 minutes. For mitochondrial stress (Fig. 2), HeLa K were treated with 75 mM NaN_3_ (Sigma) for 30 minutes at 37°C. Regarding arsenite stress cells were treated with 0.1-0.5 mM Sodium Arsenite (Sigma) for 30-60 minutes, the exact conditions are indicated for each figure in the corresponding figure legend.

Cycloheximide (Merk) was diluted in culture medium at a final concentration of 50 µg/ml. Puromycin (Sigma) was diluted in culture medium at a final concentration of 10 µg/ml.

### Creation of HeLa-UBAP2LmScarlet CRISPR cell line

HeLa K were transfected using xtremeGene9 (Sigma) with 500 ng of an all-in-one plasmid (Sigma) expressing Cas9-eGFP and the sgRNA 5’-ACAGCTGGGGGGCCAACTG-3’ (targeting UBAP2L). The all-in-one plasmids were cotransfected with 500 ng of pUC57 plasmid containing a repair template (Genewiz). The repair template was designed as a fusion of 5xGly-mScarlet flanked by two 500 bp arms, homologous to the genomic region around the Cas9 cutting site. 5 days after transfection, 3000 mScarlet positive cells were sorted in a 1 cm well and expanded for one week before a second sorting of single cells in a 96 well plates. Single colonies were grown in conditioned medium (50% fresh culturing medium and 50% filtered medium harvested from a confluent culture of HeLa K cells after 24 hours). Sorting was performed using an astrios MoFlo Cell Sorter (Beckman). After 2-3 weeks cells were screened by PCR. Briefly, cells were split and part of the cells from each clone were lysed and the DNA extracted using a commercial kit according to the manufacturer’s instructions (Quiagen). The genomic DNA obtained was subjected to PCR. Positive clones were identified according to the size of the PCR product and further screened by Western Blot. PCR amplification was performed using Taq Polymerase (Roche) according to manufacturer instruction. PCR products were separated through electrophoresis in a TBE (89 mM Tris-borate, 2mM EDTA in ddH_2_O pH 8.3) 1% Agarose gel for 20 minutes at 100 V in TBE buffer. Then the agarose gel was stained with 0.2-0.5 µg/ml Ethidium bromide in ddH_2_O for five minutes and the PCR products were visualized using an UV transilluminator (Syngene).

### Antibodies

The following antibodies were used: Mouse anti-eIF2α (1:1000 Cell Signalling L57A5), Mouse anti-G3BP (1:1000 – 1 µg/ml BD Biosciences 611126), Rabbit anti-G3BP2 (1:1000 Bethyl A302-040A) Rabbit anti-PCNA (1:1000 – 1 µg/ml Abcam ab18197), Rabbit anti-pS51-eIF2α (1:500 – 0.721 µg/ml ab32157), Mouse anti-Puromycin (1:25000 Sigma-Aldrich MABE343 [12D10]), Rabbit anti-TIA1 (1:1000 – 5 µg/ml Abcam ab40693), Mouse anti-αTubulin (1:1000 Sigma-Aldrich T9026), Rabbit anti-UBAP2L (1:2000 – 1 µg/ml Abcam ab138309), Rabbit anti-USP10 (1:200 – 1 µg/ml Abcam ab70895), AlexaFluor donkey anti-mouse 488nm (1:400 – 0.5 µg/ml Thermo Fisher A21202), AlexaFluor donkey anti-rabbit 488nm (1:400 – 4 µg/ml Thermo Fisher A11034), AlexaFluor goat anti-rabbit 594nm (1:400 – 2 µg/ml Thermo Fisher A11012), HRP-conjugated goat anti–mouse (1:10000 BioRad 170-6516), HRP-conjugated goat anti–rabbit (1:10000 BioRad 170-6515).

For U-ExM both the primary and the secondary antibody concentrations were doubled. For STED the following secondary antibodies were used: Goat anti-mouse STAR RED (1:5000 Abberior STRED), Goat anti-mouse STAR RED (1:5000 Abberior STRED), AlexaFluor donkey anti-rabbit 488nm (1:5000 – 0.2 µg/ml Thermo Fisher A11034).

### Immunofluorescence

HeLa K cells were seeded on HCl-treated coverslips and fixed with 4% formaldehyde (FA, Sigma) in Phosphate Saline Buffer (PBS, 137 mM NaCl, 2.7 mM KCl, 10 mM Na_2_HPO_4_, 1.8 mM KH_2_PO_4_ in ddH_2_O) for 10 min at room temperature (RT). Cells were permeabilized for 10 minutes with PBS-0.1% Triton X-100 (Sigma-Aldrich). For STED cells were permeabilized with PBS-0.1% Tween-20 (Sigma-Aldrich). Cells were incubated in blocking buffer (1% Bovine Serum Albumin (BSA - Sigma-Aldrich), 0.02 % NaN_3_, PBS) for 1 hr at RT and then incubated for an additional hour with primary antibodies diluted in blocking buffer. Coverslips were then washed in PBS-0.1% Tween-20 and incubated for 1 hour at RT with secondary antibodies diluted in blocking buffer and 1 mg/mL 4’6’-Diamidino-2-phenylindole (DAPI) (Sigma Aldrich). Coverslips were there mounted with either Vectashield with DAPI mounting medium (ReactoLab) or with Mowiol (Sigma Aldrich). For STED no DAPI counterstaining was performed and the samples received an additional washing step with tap water prior to mounting. The samples were mounted using ProLong™ Gold Antifade Medium (Thermofisher).

Images in Fig. 2, 3, 4 and Fig S 2 were obtained with an A1r spectral confocal microscope (Nikon) equipped with a 60× 1.4 NA CFI Plan Apochromat Lambda oil objective and 4 PMTS including 2 GaAsp for detection. Except for Fig. 1, Fig. 2 B and Fig S2 B, all the quantifications were obtained imaging the cells with either a Leica DM6000 microscope (Leica Microsystems) equipped with epifluorescence and DIC optics, 63×/1.4 numerical aperture (NA) objective, and a DFC 360 FX camera (Leica), or a Nikon microscope (Nikon) equipped with epifluorescence and DIC optics, 63×/1.4 numerical aperture (NA) objective, camera.

### Live Cell Imaging

Image series displayed in Fig. S1 F and Fig. 3 C were obtained recording cells growing in IBIDI plates at 37°C using a Nikon Eclipse Ti-E wide-field microscope (Nikon, Switzerland) equipped with a DAPI/eGFP/TRITC/Cy5 filter set (Chroma, USA), a 60× NA 1.3 oil-objective (Fig. S1 F) or a 100× NA 1.4 oil-objective (Fig. 4 C) and a Orca Flash 4.0 CMOS camera (Hamamatsu, Japan) and the NIS software. In Fig. S1 F, 11 Z-stacks (Step size = 1 µm) were taken every 60 seconds, for 30 minutes, using 5 % lamp power and recording 576 nm emission for 200 ms Exposure, with 2×2 Binning. In Fig. 4 C, 15 Z-stacks (Step size = 0.6 µm) were taken every 30 seconds, for 30 minutes, using 3% lamp power and recording 535 nm emission for 200 ms Exposure, with 2×2 Binning.

For FRAP experiment (Fig. 1 C, D and Fig S1 G, H) single plane images of HeLa UBAP2L- mScarlet were acquired at 37°C every 2 seconds using an A1r spectral confocal microscope (Nikon) equipped with a 60× 1.4 NA CFI Plan Apochromat Lambda oil objective and 4 PMTS including 2 GaAsp for detection. Imaging was performed with 2X line averaging and 0.260 µm pixel size. Photobleaching was performed with 100% 561 nm laser on a ≈2 µm diameter area (Scan Speed = 0.25). A single stress granule was followed for 10 seconds, bleached, recorded for 3 minutes (Fist FRAP cycle), bleached a second time and recorded for 3 minutes (Second FRAP Cycle). NIS Elements AR software (v.4.20.01; Nikon) was used to set acquisition parameters.

### STED super-resolution microscopy

For STED we used a pulsed gated-STED microscope DMi8 equipped with HC PL Apo 93x/ 1.30 glycerol motC STED W objective and a DFC7000T camera (Leica). Excitation was performed with a white laser, whereas depletion with either a continuous 592 nm laser or a 775 nm pulsed laser.

For 3D-STED imaging (Fig 1 A), stress granules were imaged in 3D using a 7% 635 nm laser and depletion was performed with 15% 775 nm laser, with 15% laser power in 3D STED. Imaging was performed with 4X line averaging and ≈20 nm pixel size. 3D stacks were acquired with a step size of 100 nm. For 2D-STED dual color (Fig. 1 E) stress granules were imaged in 2D using a 7% 635 nm excitation laser + 15% 775 nm depletion laser, followed by a 10% 488 nm excitation laser + 15% 592 nm depletion laser. Imaging was performed with 4X line averaging and ≈20 nm pixel size. Images were deconvolved using Huygens (Scientific Volume Imaging).

### Ultrastructure Expansion Microscopy (U-ExM)

U-ExM was performed as described in (Gambarotto et al., 2019). Briefly, in Fig. 1 G, H, I the cells were grown in 12 mm slides and then fixed with 4% FA (Sigma) in PBS for 10 min at 37°C. Then, coverslips were incubated in a solution of 0.7% FA, 1% AA (Acrylamide, Sigma) in PBS for 5 h at 37 °C. Gelation was performed in a precooled humid chamber with coverslips facing up with 35 μl of monomer solution (19% Sodium Acrylate (SA, Sigma), 10% AA, 0.1% BIS (N,N’-methylenbisacrylamide, Sigma) in PBS supplemented with 0.5% APS and 0.5% TEMED. Immediately after the addition of MS the chamber was closed with a second coverslip on top. Coverslips were then incubated for 1 hour at 37 °C in the dark. Coverslips with gels were transferred into 2 ml of denaturation buffer (200 mM SDS (Sodium dodecyl sulfate, Sigma), 200 mM NaCl (Sigma), and 50 mM Tris (Biosolve) in ddH_2_O, pH 9) for 15 min at RT. Samples were then placed into a 1.5-ml Eppendorfs filled with fresh denaturation buffer, and incubated at 95 °C for 1 hour. After denaturation, gels were expanded in beakers filled with ddH_2_O. Water was exchanged three times every 30 min at RT, for the last wash samples were incubated O/N in ddH_2_O at 4°C. Next, gels were placed in PBS two times for 15 min. Primary antibodies were diluted in 2% BSA PBS and binding was performed at 37 °C for 3 h. Gels were then washed in 0.1% Tween 20 PBS three times for 10 min at RT. The same steps were repeated for secondary antibodies. Finally, gels were placed in ddH_2_O. Water was exchanged three times every 30 min, and then gels were incubated in ddH_2_O overnight.

For imaging, a piece of gel was mounted in a 2-well IBIDI plate coated with poly-L-lysine (Sigma). Confocal microscopy was performed using an A1r spectral confocal microscope (Nikon) equipped with a 60× 1.4 NA CFI Plan Apochromat Lambda oil objective and 4 PMTS including 2 GaAsp for detection. 3D z-stacks at 100 nm steps were acquired with a pixel size of 50 nm. Images were deconvolved using Huygens.

### Western Blotting

HeLa extracts were prepared from HeLa K or HeLa UBAP2L-mScarlet cells by scraping the cells in 50 mM Tris-HCl (pH 7.4), 150 mM NaCl, 1 mM EDTA, 1 mM EGTA, 0.27 M sucrose, 1% Triton X-100, complete protease inhibitor cocktails (Roche), and phosphatase inhibitor PhosSTOP (Roche) for 1 hour on ice before clarification by centrifugation (14’000 g 20 minutes, 4°C). Supernatants were collected and quantified using Bradford Reagent (Bio-Rad Laboratories). Lysates were then diluted in Laemmli buffer and a 30 µg of lysates were separated by SDS-PAGE using a manually poured 10% Acrylamide Gel. Proteins were then transferred onto a nitrocellulose membrane (Sigma). The membrane was blocked with 5% BSA in TBS (Tris-Buffered Saline, 50 mM Tris-Cl, 150 mM NaCl, in ddH_2_O, pH 7.4) and incubated overnight with primary antibodies at 4°C in the same solution. The following day the membrane was washed with 0,1% Tween TBS and incubated with secondary antibodies in 0,1% Tween TBS. Proteins were visualized with ECL (Millipore).

Western blot quantification (Fig S2 C) was performed calculating the area under the curve using Fiji’s “Gel” plugin. UBAP2L quantification was normalized on the corresponding tubulin signal and then on the average UBAP2L levels in control depleted cells of 4 independent experiments. Resulting values were scaled from 0 to 100. Statistical analysis was performed with Prism 8 (GraphPad).

### Particle Diameter Measurement, Line Profiles and stress granule areas

In Fig. 1 B and Fig. S1 A particle diameters was measured using Imaris 9.2 (Bitplane) and data were analyzed with Python 3.7.0 and the libraries “pandas”, “matplotlib.pyplot” and “numpy”. Particles below 0.06 µm were excluded from the analysis (8.75 % of the G3BP1 particles, 4.87 % of the UBAP2L particles). n= 857 particles (44 stress granules, 38 cells) for G3BP, n= 904 particles (84 stress granules, 67 cells) for UBAP2L.

In Fig. 1 F, I line profiles were calculated for both channels drawing a line on a stress granule and using FiJi “Plot Profile” function. The pixel by pixel gray value was normalized on the average grey value of all pixels in the line. Data were analyzed with Prism 8. In Fig. 1 G the 3D surface of the stress granules particles was computed using Imaris 9.2.

To quantify stress granule areas in Fig. 3 E, F we created a Image J macro available on github: https://github.com/LCirillo/FiJiMacro/blob/master/IJ_Macro_SG_Area. For each independent experiment a minimum of 53 stress granules and were measured for each condition.

### FRAP quantification

In Fig. 1D and Fig. S1 G, H, the bleached stress granule was tracked using Imaris 9.2 and the mean grey intensity over time was measured (FI). In parallel, the total fluorescence of the image (FI_tot_). After background subtraction the data were normalized as follows to account for photobleaching and fluorophores loss:

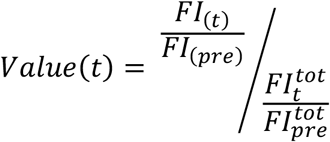

All the particles that did not recover to 1 in the second FRAP were excluded from the final analysis (32.5 %). Data analysis was carried out using Prism 8. Result were averaged separately for the 1st FRAP and the 2nd FRAP and plotted as mean +/- 95% CI. The mean recovery curve was fitted to a single exponential using Fiji (FRAP1 R2= 0.985, FRAP2 R2=0.988) and the immobile fraction extrapolated. Statistical analysis was performed with Prism 8.

### Colocalization analysis

In 2D STED and Expansion microscopy (Fig. 1 E, H) Pearson’s Correlation Coefficient was measured using Imaris 9.2. n= 20 stress granules from 11 cells (2D STED), n=28 stress granules from 7 cells (U-ExM).

For the cross-correlation analysis of the distributions of UBAP2L and G3BP1 (Fig. 2 B, Fig. S2 B) the Pearson’s Correlation Coefficient was measured using single plane confocal images of HeLa K cells experiencing different stresses. The Pearson Coefficient was measured using the Fiji plugin “Coloc2”. Cololcalization analysis was performed as previously described (Martino et al., 2017). Statistical analysis was performed using Prism 8.

### Polysome Decay Assay

To segment and quantify polysome decay in Fig. 4 C we created an Image J macro available on github: https://github.com/LCirillo/FiJiMacro/blob/master/IJ_Macro_PolysomeCounting.

The number of polysomes obtained was divided by the surface occupied by the cells to obtain the polysome density (PD). The polysome density at each time point (Pd_t_) was normalized by the polysome density at the first frame of imaging (Pd_0_) to obtain the normalized polysome density.

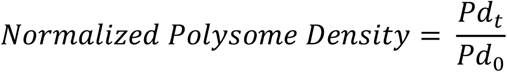

Data analysis was carried out using Prism 8.

### Figure assembly

All the figures were assembled using Adobe Illustrator CS6.

## ACKNOWLEDGMENTS

We are grateful to D. Gambarotto and P. Guichard for the U-ExM protocol. We thank N. Kedersha for the G3BP1 and 2 Knock out cell lines. We thank X. Pichon and E. Bertrand for sharing the polysome reporter cells line. We thank the Flow Cytometry facility and the Bioimaging Core facility of the Medical Faculty, in particular F. Prodon for his help with the STED nanoscopy and N. Liaudet for advices in image processing and analysis. We are grateful to all members of the laboratories of Monica Gotta and Patrick Meraldi for fruitful discussions.

This work was supported by a grant from the Swiss National Science Foundation and by funding from the University of Geneva to M. Gotta.

## AUTHOR CONTRIBUTIONS

A.C. and L.C. performed all the experiments, quantifications and interpreted the results. L.C. and M.G wrote the manuscript and conceived the study. M.G. acquired funding.

---

**Figure Supplementary 1.**
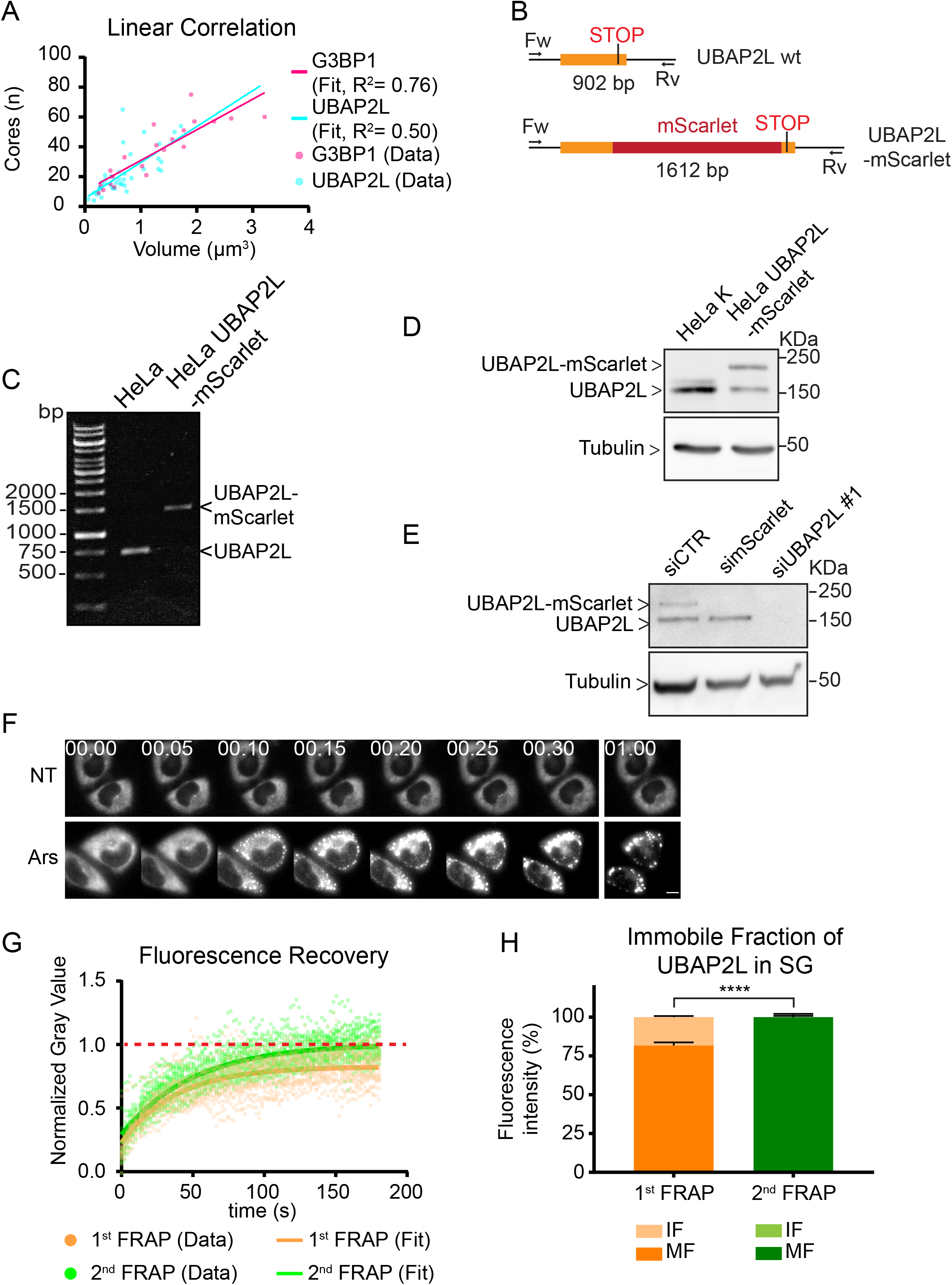
Development of a UBAP2L-mScarlet HeLa cell line. A) Linear regression of stress granule size measured by 3D-STED versus the number of particles per granule. B) Schematic representation and screening strategy used to identify UBAP2L-mScarlet fusion clones. C) Representative image of PCR products of parental HeLa cells compared to a clone with successful mScarlet insertion, as indicated. D) Image of a Western blot of lysates from HeLa K cells (left) compared to HeLa UBAP2L-mScarlet cells (right). In both (D) and (E) the membrane was blotted with anti-UBAP2L (upper panel), and anti-Tubulin (lower panel) antibodies. E) Image of a Western blot of lysates from HeLa UBAP2L-mScarlet cells treated with different siRNAs. F) Maximum projections of wide-field images from time-lapse movies of HeLa UBAP2L-mScarlet cells non treated (NT) and treated with 0.5 mM arsenite. Time is expressed at hh:mm, top left of each image. Scale bar corresponds to 10 μm. G) Graph showing the FRAP recovery of the first FRAP cycle and the second FRAP cycle of all the cells analyzed in Fig. 1 D. The continuous lines represent the single exponential functions used to fit the mean of the recovery curves. The red dotted line centered at 1 highlights complete fluorescence recovery. H) Histogram representing the percentage of mobile fraction (MF) and immobile fraction (IF) in the two FRAP cycles. Error bars indicate SD. Statistical significance was determined using ANOVA and Dunnett’s multiple comparison.

**Figure Supplementary 2.**
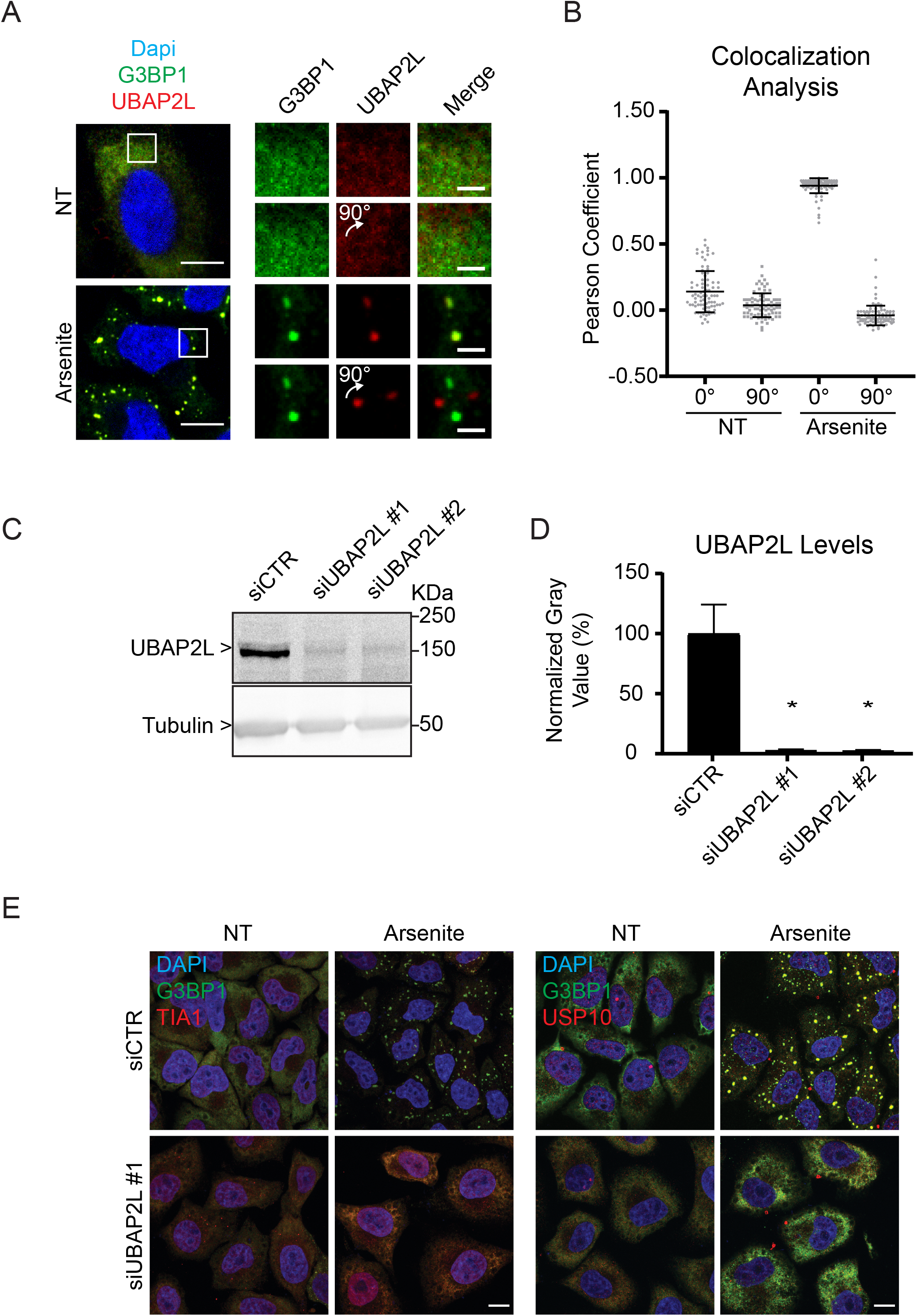
UBAP2 colocalizes with G3BP1 in arsenite-induced stress granules and it is required for stress granules assembly. A) Left: Confocal images of HeLa K cells stained with anti-G3BP and anti-UBAP2L antibodies and counterstained with DAPI. Cells were either non treated (first row) or treated with sodium arsenite 0.1 mM for 30 min (second row). Scale bar corresponds to 10 μm. Right: magnification of a ROI of the images on the left (white box). Scale bars correspond to 1 μm. B) Cross correlation between G3BP and UBAP2L signal before and after a 90° rotation of the image. Each dot corresponds to the Pearson coefficient of a single ROI. Error bars indicate SD. 82 (Control) and 100 (arsenite) cytoplasmic ROIs from 3 independent experiments were analyzed. C) Representative image of a Western blot of lysates from HeLa K cells transfected with a control siRNA (first lane) compared to lysates of HeLa K transfected with a siRNAs targeting UBAP2L (2^nd^ and 3^rd^ lane). The membrane was blotted with anti-UBAP2L and anti-αTubulin, as indicated. D) Percentage of UBAP2L depletion in HeLa K cells treated with the indicated siRNAs, quantified from western blots. Error bars indicate Standard Error of the Mean (SEM). Statistical significance was determined using ANOVA and Dunnett’s multiple comparison. N=4 independent experiments. E) Maximum projections of confocal images of HeLa K cells transfected with the indicated siRNAs and stained with different stress granule markers. Cells treated for 30 minutes with 0.1 mM arsenite or left untreated. Left: anti-G3BP and anti-TIA1 antibodies and counterstained with DAPI. Right: anti-G3BP and anti-USP10 antibodies and counterstained with DAPI. Scale bars correspond to 10 µm.

**Figure Supplementary 3.**
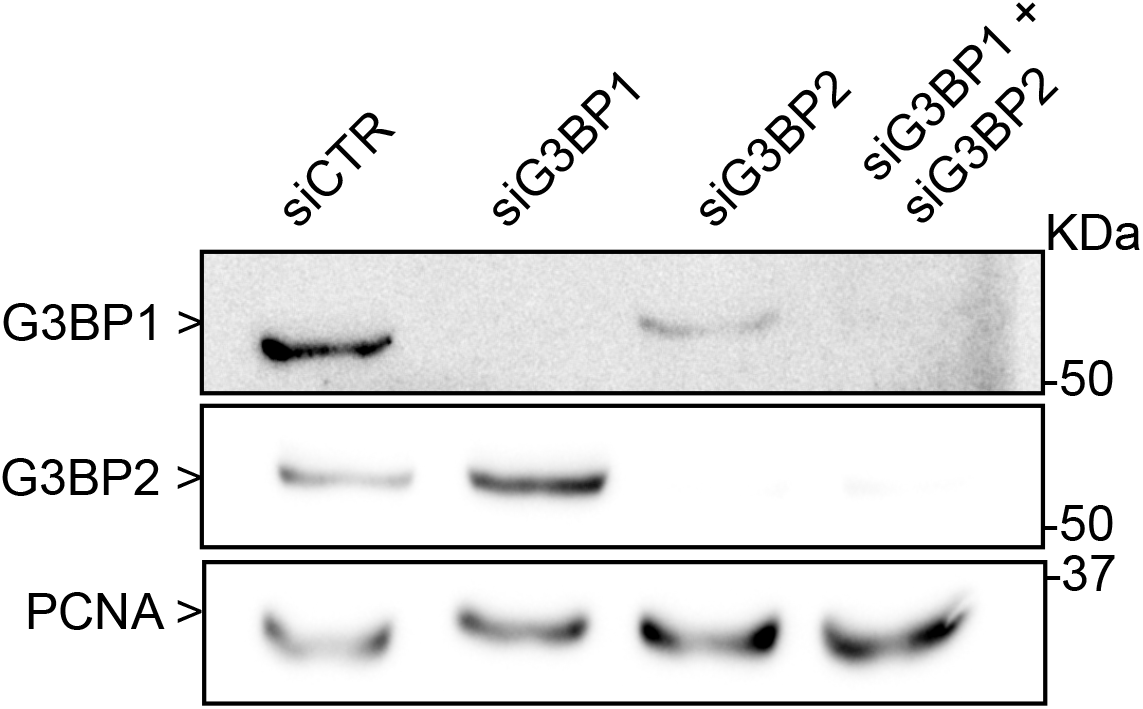
Depletion of G3BP1 and 2. Image of a Western blot of lysates from HeLa K cells transfected with a control siRNA (1^st^ lane) compared to lysates of HeLa K transfected with a siRNAs targeting G3BP1 and 2 (2^nd^, 3^rd^ and 4^th^ lane). The membrane was blotted with anti-G3BP, anti-G3BP2 and anti-PCNA (loading control) antibodies as indicated.

